# Structural basis for tunable affinity and specificity of LxCxE-dependent protein interactions with the retinoblastoma protein family

**DOI:** 10.1101/2022.06.10.495386

**Authors:** Sivasankar Putta, Lucia Alvarez, Stephan Lüdtke, Peter Sehr, Gerd A. Müller, Samantha M. Fernandez, Sarvind Tripathi, Joe Lewis, Toby J. Gibson, Lucia B. Chemes, Seth M. Rubin

## Abstract

The retinoblastoma protein (Rb) and its homologs p107 and p130 are critical regulators of gene expression during the cell cycle. Functional inactivation of Rb family proteins leads to loss of cell-cycle control and promotes genome instability and proliferation, which are hallmarks of cancer. Rb proteins share a structural domain, known as the “pocket domain”, which mediates association with a large number of cellular proteins. A cleft in the pocket domain binds an LxCxE sequence motif in these proteins, many of which function with Rb proteins to co-regulate transcription during quiescence and G_1_. Proteins from oncogenic DNA viruses also bind this cleft to inactivate Rb family proteins, and the E7 protein from the human papillomavirus has been used as the primary model for understanding LxCxE motif interactions. Evidence presented previously and here demonstrates that the E7 sequence binds tighter and competes with cellular proteins for the LxCxE site, but the molecular basis for the relatively weak affinities of the cellular proteins has been poorly understood. Here, we present binding data and a series of crystal structures of the p107 pocket domain in complex with LxCxE peptides derived from the transcriptional co-repressor proteins HDAC1, ARI4DA, and EID1. The structures reveal determinants of protein binding beyond the L-C-E core that include the N and C-terminal sides of the LxCxE motif and the x (wild card) residues. Our results explain how differences in these interactions outside the L-C-E core result in weaker affinities of pocket protein for cellular LxCxE proteins compared to E7 and result in differences in Rb and p107 affinities for some LxCxE-containing sequences. Our study provides new insights into how Rb proteins bind their myriad of cell partners and lays the foundation for structure-based design of peptides that inhibit oncogenic viral protein interactions.

## Introduction

The Rb tumor suppressor protein is critical for normal development and is commonly found inactivated in cancer (Sherr and McCormick, 2002). Rb has numerous activities centered around regulation of cell-cycle dependent gene expression and genome maintenance (Burkhart and Sage, 2008; Dick and Rubin, 2013). In addition to its best-studied function of inhibiting E2F transcription factors, Rb also plays a role in genomic repeat silencing, cell lineage stability, DNA break repair, DNA replication, and formation of higher-order chromatin structures (Dick et al., 2018). Many of these functions are mediated by yet poorly understood protein interactions. Cells that lack Rb or contain Rb protein that is constitutively inactivated by upstream kinases or DNA tumor viral proteins are prone to genotoxic stress and transformation, and therapeutics that restore Rb activity are now being used to treat cancer (Knudsen et al., 2019).

Many Rb functions, for example negative regulation of cell-cycle progression and inhibition of E2F dependent transcription, are shared by two close paralogs p107 and p130 (van den Heuvel and Dyson, 2008). Rb family proteins repress gene expression during quiescence and the G1-S transition by binding E2Fs, which inhibits E2F transactivation, and recruiting co-repressor proteins like HDAC1 (Histone deacetylase 1), EID1 (EP300-interacting inhibitor of differentiation 1), and ARID4A (AT-rich interaction domain-containing protein 4a) to E2F promoters (Dick and Rubin, 2013; Ferreira et al., 1998; Lai et al., 1999; MacLellan et al., 2000; Miyake et al., 2000). Despite these functional redundancies, Rb is found more commonly mutated in human cancer than p107 and p130, and loss of Rb is sufficient to promote tumorigenesis in mouse models (Classon and Harlow, 2002; Knudsen et al., 2019; Mulligan and Jacks, 1998). The molecular differences between Rb and its paralogs that account for the more potent tumor suppressor capacity of Rb are still incompletely understood. There is evidence that interactions with E2Fs vary (Liban et al., 2017; Liban et al., 2016; Trimarchi and Lees, 2002), and interactions with co-repressors have been better characterized for Rb, although it has been reported that p107 binds ARID4A and HDACs (Ferreira et al., 1998; Lai et al., 1999).

The Rb family proteins act as scaffolds that assemble diverse protein complexes in a manner that is regulated by multisite phosphorylation (Dick and Rubin, 2013; Morris and Dyson, 2001; Rubin, 2013). Therefore, determining molecular details for how Rb proteins interact with cellular partners is critical for understanding function and regulation of these important tumor suppressors. Rb proteins contain three primary functional domains: an N-terminal domain, a central “pocket” domain, and an intrinsically disordered C-terminal domain (Figure 1A) (Dick and Rubin, 2013). The loops that connect these domains and phosphorylation sites in the proteins mediate different conformations that control binding of more than 200 proteins (Morris and Dyson, 2001; Rubin, 2013; Sanidas et al., 2019). The structured pocket and N-terminal domains, which are well conserved in the family, each contain two tandem cyclin folds (A and B subdomains) that are connected by a disordered linker (Hassler et al., 2007; Lee et al., 1998). The pocket domain is necessary for nearly all Rb protein functions, including inhibition of E2F-dependent transcription (Dick and Rubin, 2013). The E2F transactivation domain binds at the interface between the A and B sub-domains (Lee et al., 2002; Xiao et al., 2003). The B subdomain contains the site that binds an LxCxE sequence motif present in many cellular and viral interacting proteins (Lee et al., 1998). Many functions outside of E2F association are mediated through the LxCxE cleft of the pocket proteins (Dick et al., 2018; Dick and Rubin, 2013). For example, the LxCxE-binding cleft is required for regulation of heterochromatin structures and proper mitotic condensation through recruitment of the condensin II complex (Coschi et al., 2010; Manning et al., 2010).

**Figure 1.**
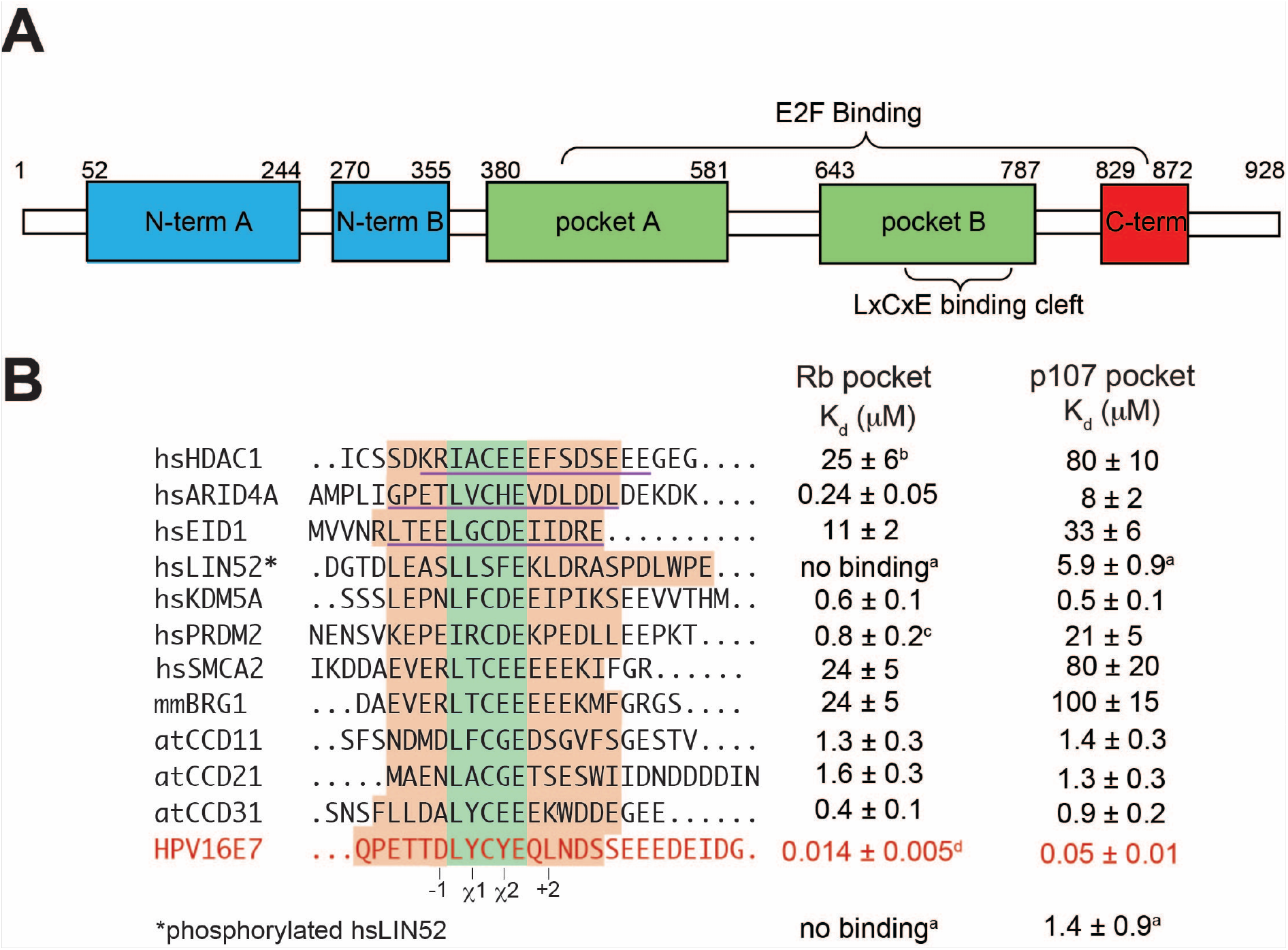
Affinity measurements of LxCxE peptides for Rb and p107. (A) Schematic diagram of the domain organization of Rb. A similar domain architecture is observed in p107 and p130. (B) Alpha Screen affinity measurements of cellular LxCxE peptides and comparison to the viral E7 peptide. In the sequence alignment, the peptide sequences used for affinity studies are colored peach, the core LxCxE motif is colored green, and the peptide sequence used in structural studies is underlined. The position of variable amino acids relative to the core (−1, +2) and the wild-card positions in the core motif (χ1 and χ2) are labeled. hs is Homo sapiens, mm is Mus musculus, and at is Arabidopsis thaliana. ^a^Reported in (Guiley et al., 2015; Singh et al., 2005) ^b^Measured here and also reported as K_d_ = 0.64 ± 0.07 μM in (Sun et al., 2015). ^c^Measured here and also reported as K_d_ = 5 ± 1 nM in (Chemes et al., 2011). ^d^Measured here and also reported as K_d_ = 10 ± 3 μM in (Singh et al., 2005).

The LxCxE-cleft in the pocket domain is also used by Rb family proteins to regulate gene expression at E2F promoters. Interaction between an LxCxE-like motif in the protein LIN52 and p107 and p130 mediates formation of the DREAM complex, which represses cell cycle genes during quiescence (Guiley et al., 2015; Litovchick et al., 2011; Litovchick et al., 2007). In addition, transcriptional co-repressor proteins like HDAC1, ARID4, and EID1 contain the LxCxE motif (Brehm et al., 1998; Defeo-Jones et al., 1991; Fattaey et al., 1993; Ferreira et al., 1998; Luo et al., 1998; MacLellan et al., 2000; Magnaghi-Jaulin et al., 1998; Miyake et al., 2000). Pocket proteins recruit the histone deacetylase HDAC1 to inhibit E2F-dependent transcription by modulating chromatin structure and the accessibility of co-activators and transcriptional machinery (Ferreira et al., 1998; Zhang et al., 2000). ARID4A (also known as Rb-binding protein 1 or RBP1) blocks E2F mediated gene transcription by forming complexes with Rb proteins, E2F, and HDAC1 at promoter sites and remodeling chromatin (Lai et al., 1999). The co-repressor EID1, which plays a role in promoting differentiation, binds to the Rb LxCxE cleft, and inhibits the histone acetyltransferase (HAT) activity of the co-activator protein p300 (MacLellan et al., 2000; Miyake et al., 2000). Although genetic ablation of the LxCxE-cleft in Rb results in normal cell growth and animal development, it leads to changes in chromatin structure, elevated cell-cycle gene expression, and increased cancer susceptibility in response to DNA damaging agents (Bourgo et al., 2011; Gonzalo et al., 2005; Talluri and Dick, 2012). These reports and bioinformatics analysis suggest the involvement of the LxCxE-cleft in binding a large set of proteins required for the regulation of cell functions (Kumar et al., 2021; Palopoli et al., 2018). While many cellular proteins are known to use LxCxE motifs to bind to pocket proteins, the molecular basis of these interactions is, for the most part, unknown.

In addition to the cellular proteins, several viral oncoproteins, including adenovirus E1A, SV40 T antigen (TAg), and human papillomavirus E7, use LxCxE motifs to bind and suppress the function of Rb family proteins (DeCaprio, 2009; Kim et al., 2001; Lee et al., 1998; Palopoli et al., 2018). These motifs provide a high affinity contact to Rb proteins (Chemes et al., 2010; Jones et al., 1990; Lee et al., 1998; Singh et al., 2005), which in conjunction with a second binding domain in the viral protein that directly competes with E2F, disrupts Rb protein-E2F complexes. Without the LxCxE motifs, viral proteins lose their capacity to bind and to deregulate the Rb function in cell cycle. Structural studies of E7 bound to Rb and p107, TAg bound to Rb, and the cellular protein LIN52 bound to p107 have revealed important details about the LxCxE-binding mode, including the key interaction made by the core leucine, cysteine, and glutamate residues (Guiley et al., 2015; Kim et al., 2001; Lee et al., 1998). In the case of LIN52, the central cysteine is replaced by a serine, and the resulting loss of affinity is compensated for by interactions with a phosphoserine that is C-terminal to the core motif. This sequence design creates a phosphoswitch that allows the interaction to be regulated by an upstream kinase and also makes association specific for p107/p130 over Rb (Guiley et al., 2015).

These observations regarding LIN52 association raise the questions of whether cellular LxCxE motifs generally bind Rb proteins with much weaker affinity than E7 and other viral proteins and whether the relatively weak affinity is important for regulation. Differences in affinity between different LxCxE-containing proteins and the modulation of Rb binding by mutations to regions flanking the core motif also suggest that interactions beyond the core motif are critical for fine tuning of the LxCxE affinity (Jones et al., 1990; Palopoli et al., 2018; Singh et al., 2005). This idea is further motived by a growing body of evidence indicating that linear motif binding-affinity and specificity determinants are encoded by contextual features that include the wildcard positions, the residues flanking the core motif, and the conformation of the intrinsically disordered regions harboring the motifs (Bugge et al., 2020; Prestel et al., 2019). In the case of the LxCxE motif interaction with pocket proteins, limited structural knowledge, particularly of how cellular proteins bind, has limited our understanding of the key binding determinants. Here we present biophysical evidence that interactions between a set of cellular LxCxE sequences, originally identified as Rb-binders, and pocket domains are weaker than interactions with the viral E7 motif and that these sequences tend to bind Rb with higher affinity than p107. We report high-resolution structures of cellular HDAC1, ARI4DA and EID1 LxCxE motif peptides bound to the human p107 pocket domain. The structures reveal additional determinants of peptide binding at the LxCxE site that include interactions N- and C-terminal to the core motif residues and the importance of bulky residues in the x “wild-card” positions. An engineered HDAC1 protein that contains an artificially high affinity LxCxE sequence binds robustly to Rb in cells. These data reveal the mechanism for a fine-tuned interaction between Rb proteins that is balanced for affinity and regulation, and they could guide the design of inhibitors to prevent Rb inactivation by the viral oncoproteins.

## Results

### Cellular protein LxCxE sequences bind Rb tighter than p107 and bind both pocket domains with weaker affinity than the prototypical HPV16 E7 viral protein

Previous measurements of binding affinities between LxCxE-containing peptides from HDAC1 and LIN52 for the Rb and p107 pocket domain demonstrate K_d_ values on the order of ~10 μM (Guiley et al., 2015; Singh et al., 2005), while the IxCxE motif from PRDM2 binds to Rb with ~0.6 μM affinity (Sun et al., 2015). These affinities are up to several hundred-fold weaker than for the LxCxE motif from the viral HPV16 E7 protein. In order to compare a larger sample of cellular proteins, we performed an Alpha Screen assay to measure binding constants of several synthetic peptides containing LxCxE sequences from known Rb family interacting proteins (Figures 1B and S1 and Table S1). We focused on transcriptional co-repressor proteins (ARI4DA, KDM5A, SMCA2, PRDM2, HDAC1, EID1, and BRG1) and proteins that regulate Rb phosphorylation (CycD paralogs from *Arabodopsis thalania* were used for improved peptide synthesis), and we measured affinities for both the Rb and p107 pocket domains. The results obtained with the Alpha Screen assay for HDAC, PRDM2 and HPV16 E7 were in very good agreement with the previously measured affinities (Figure 1B) (Chemes et al., 2011; Singh et al., 2005; Sun et al., 2015).

Our results suggest that cellular LxCxE sequences generally bind weaker to pocket proteins than the HPV16 E7 peptide and bind with a wide range of affinities from K_d_ ~250 nM to ~100 μM. For many of the co-repressor proteins, binding of the peptides to the Rb pocket domain occurred with higher affinity than to the p107 pocket domain. We observe affinity differences up to 30-fold in the cases of ARID4A and PRDM2, whereas affinities are similar for all the CycD peptides and KDM5A. This observation is notable considering that the residues in the pocket cleft that contact the core residues in the LxCxE motif (LCE) are identical across Rb family proteins with a few exceptions that we explore below (Guiley et al., 2015; Lee et al., 1998). Overall, the affinity measurements performed here support the conclusion that subtle sequence differences outside the core LCE residues can greatly tune binding affinity. This conclusion is also supported by previous mutational studies and sequence analysis, which revealed common sequence determinants that lie outside of the core residues. These include residues in the “χ” or “wild-card” positions, acidic residues in positions −1 to −3 from the core L residue, a stretch of acidic residues and/or phosphorylation sites located C-terminal to the core LxCxE motif, and a conserved hydrophobic residue located C-terminal to the LxCxE motif, often at the +2 position (Figure 1B) (Chemes et al., 2011; Chemes et al., 2010; Guiley et al., 2015; Jones et al., 1990; Kumar et al., 2021; Palopoli et al., 2018; Singh et al., 2005).

### Crystal structures of the p107 pocket domain in complex with LxCxE peptides from transcriptional co-repressors

In order to understand the structural basis for how subtle sequence differences can influence LxCxE-protein affinity, we conducted X-ray crystallography studies on LxCxE peptides from cellular proteins in complex with the pocket domain. We selected the minimal regions of several cellular LxCxE motif peptides based on their solubility and likely potential for crystallization (Figure 1B, underlines sequences). For the pocket domain we used a protein construct p107ΔL (residues 389–972, Δ600-779, Δ888-923), which contains the human p107 pocket domain sequence but lacks the loop between the two structured subdomains and a loop within the B subdomain. We were previously successful in solving structures of E7 and LIN52 peptides bound to the p107 pocket domain using this construct (Guiley et al., 2015), and we found we could reproducibly grow crystals in similar conditions with several LxCxE peptides. Crystals were grown of p107ΔL pocket domain with human HDAC1, ARID4A, and EID1 LxCxE peptides that diffracted to 2.6 Å, 2.7 Å, and 2.2 Å respectively (Table S2 and Figure 2). Structures were determined by molecular replacement using the p107 pocket domain from the E7 complex structure (PDB: 4YOZ) as a search model. When bound to the HDAC1 or EID1 peptides, the p107 domain crystallized with one complex in the asymmetric unit in the C2_1_ space group, while the ARID4A-p107ΔL complex crystallized with two copies in the P1 space group. As expected, all three peptides bind to the B subdomain at a site distinct from where the E2F transactivation domain binds (Figure 2B-2D). The pocket domain shows minimal differences among the structures and in comparison to the domain bound to the E7 LxCxE peptide (Figure S2).

**Figure 2.**
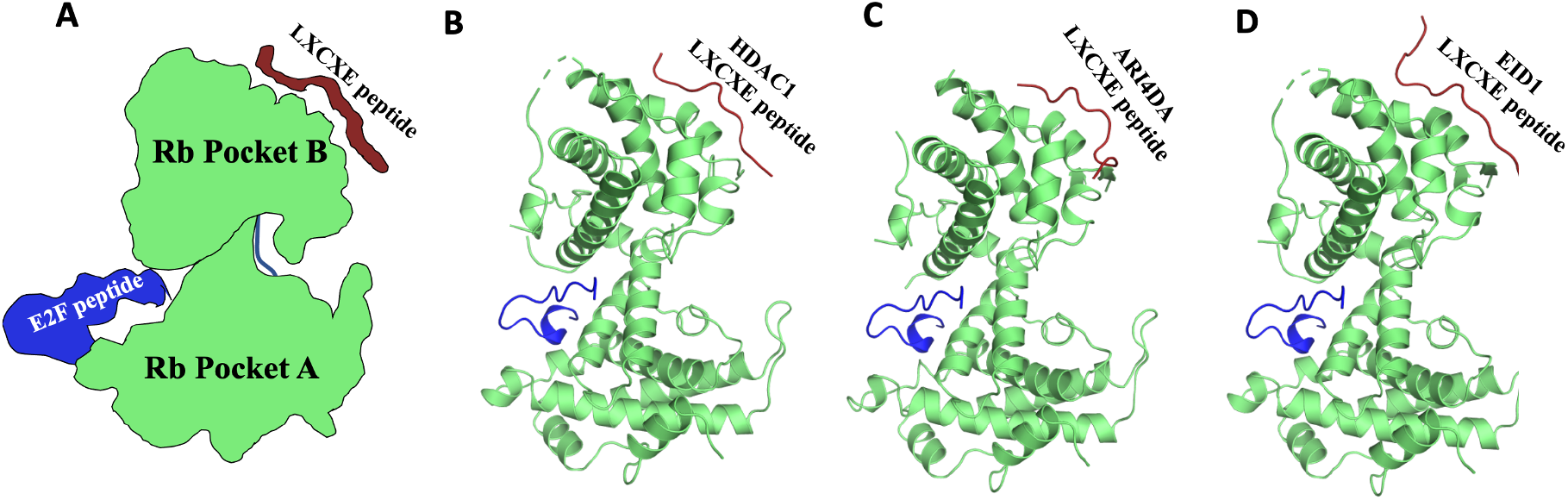
Structures of P107ΔL with cellular LxCxE peptides. (A) Schematic of the architecture of the p107 pocket domain showing the E2F and LxCxE binding sites. (B, C, and D) Ribbon models of p107ΔL pocket domain with human HDAC1, ARI4DA, and EID1 LxCxE motif peptides. The pocket domain shown in green, and the E2F and LxCxE peptides are shown in blue and red, respectively. The E2F peptide shown here is from PDB: 1N4M is superposed to show the different locations of the binding sites.

### Cellular LxCxE peptide interactions beyond the core motif

The LxCxE peptides from the transcriptional co-repressor proteins all display similar interactions made by the alternating LCE residues as previously observed for the viral E7 and TAg sequences (Guiley et al., 2015; Kim et al., 2001; Lee et al., 1998) (Figure 3). Briefly, the leucine and cysteine insert into shallow hydrophobic pockets formed between the third and fifth helices of the B subdomain cyclin fold, and the glutamate sidechain makes backbone hydrogen bonds that cap the N-terminus of the fourth helix in the fold. While these interactions are invariant, we observe a number of interactions and structural features that vary among the co-repressor protein peptides and are different from the mode of binding of the tighter affinity viral peptides. These include interactions N-terminal and C-terminal to the core LxCxE motif and the position and interactions of the wild-card sidechains.

**Figure 3.**
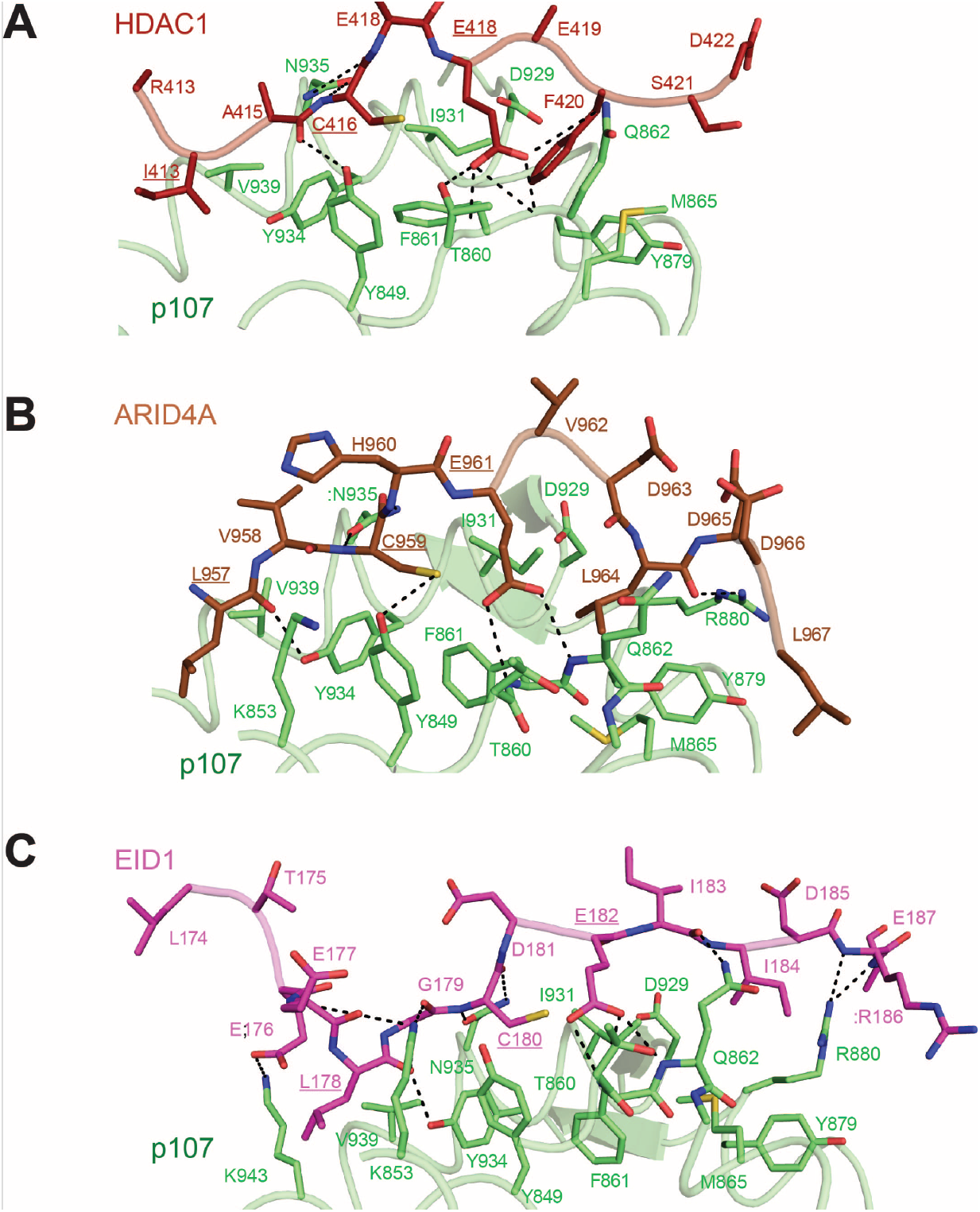
Comparison of binding interactions made by cellular LxCxE peptides. p107ΔL pocket domain in complex with peptides from (A) HDAC1, (B) ARI4DA, and (C) EID1. The core LCE motif residues are underlined.

#### Interactions N-terminal to the LxCxE motif

The viral E7 protein has an acidic aspartate at the −1 position relative to the start of the LxCxE motif, and an acidic residue is present in position −1 to −3 in all the other cellular peptides in our affinity study (Figure 1B). In some of the low affinity cellular peptides (e.g. HDAC1, SMCA2, and BRG1) there are also positive charges N-terminal to the LxCxE motif. It was previously reported that replacing the N-terminal basic arginine residue at the −1 position in the HDAC peptide for an acidic aspartate residue (R413D) increased the affinity 2-fold, suggesting that interactions involving acidic residues at the −1 position contribute mildly to stability (Singh et al., 2005). In the Rb-bound E7 (1GUX) structure, the acidic aspartate residue in the −1 position (D21) makes a long-distance hydrogen bond interaction with the χ2 sidechain (Y25) of the E7 peptide (Figure 4A); however, the same D21 points away from Y25 in the structure of E7 bound to p107 (Figure 4B). The −1 position of the SV40 TAg LxCxE motif points away from the binding groove, but it is part of an alpha helix structure (Figure 4C). The cellular peptide we studied that contains a −1 acidic is EID1 (E177), and the structure shows a novel interaction made at this position. E177 makes a salt bridge with K853, and K853 is repositioned relative to the other structures such that an additional hydrogen bond is made with the E177 backbone carbonyl (Figure 4D). Taken together, these results suggest that the positioning and interactions made by the acidic residues at the N-terminus of the motif can vary.

**Figure 4.**
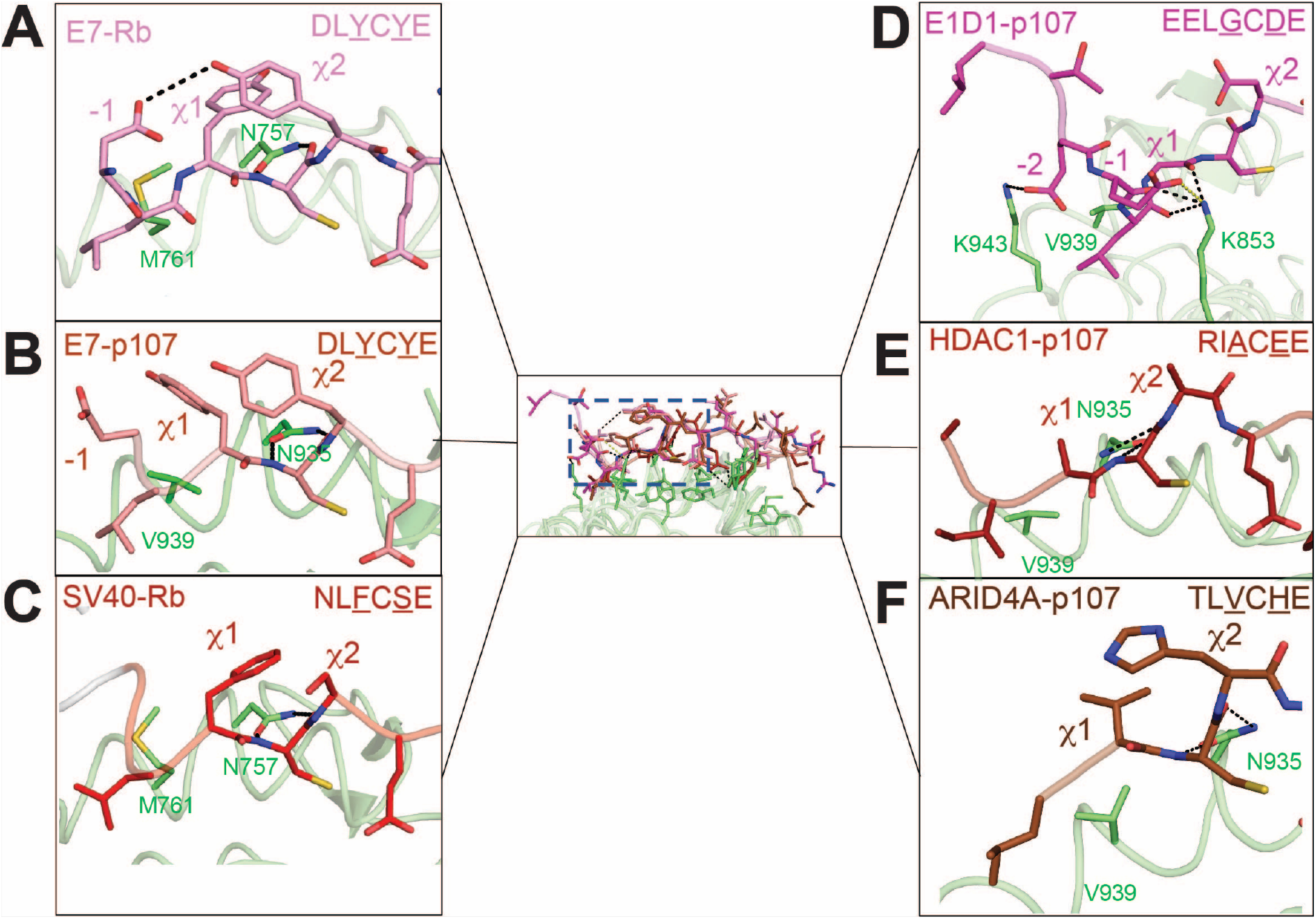
Interactions N-terminal to the core LXCXE motif and position of wild-card residues. The aligned structures of (A) E7-Rb, (B) E7-p107, (C) SV40-Rb, (D) EID1-p107, (E) HDAC1-p107, and (F) ARI4DA-p107. Peptides are shown overlaid in the middle panel and are separated in the peripheral panels to show the individual conformations of the wild-card sidechains (χ1 and χ2, underlined in seuence) and N-terminal (−1, −2, etc.) acidic residues.

The p107-bound EID1 structure also reveals an interaction made by an acidic residue in the −2 position (E176), which forms a salt bridge with K943 in p107 (Figure 3C and 4D). It is not clear why EID1 has a uniquely ordered N-terminus. To assess whether the structured acidic residues located at positions −1 and −2 of EID1 contributed stabilizing interactions, we measured by isothermal titration calorimetry (ITC) the binding affinity of an EID1 peptide in which we mutated the −1 and −2 glutamates to arginine (E176R/E177R). This mutation resulted in a subtle loss of affinity for Rb (K_d_ = 6 ± 3 μM for WT and K_d_ = 9 ± 3 μM for mutant) and p107 (K_d_ = 7 ± 5 μM for WT and K_d_ = 16 ± 5 μM for mutant) (Figures S3A-S3D), which suggested these acidic residues contribute mildly to complex stability. An acidic residue is present in position −2 in ARID4A and in position −3 in LIN52, for which there are now structures available. However, the N-terminal acidic residues are not visible in the electron density in those structures. We could only build the ARID4A peptide in the structure solved here starting at L957, which is the canonical Leu in the motif (Figure 3B). The stretch of acidic residues C-terminal to the LxCxE motif in E7 plays a stabilizing role by establishing long-range electrostatic interactions with the positively charged residues surrounding the LxCxE cleft (Chemes et al., 2011; Chemes et al., 2010). The fact that N-terminal acidic residues may remain unstructured suggests that these residues could play a stabilizing role similar to the C-terminal acidic stretch through dynamic and more diffuse electrostatic interactions with the positive residues near this side of the cleft, especially K853 and K943 (K713 and K765 in Rb), rather than making specific ordered contacts as observed in EID1. These dynamic interactions could confer different degrees of affinity enhancement, helping to fine-tune the binding affinity of the core LxCxE motif.

#### Wild-card residues of the LxCxE motif

Based on the E7 structure and mutagenesis data, it has been suggested that the affinity of LxCxE-sequences depends on the identity of the residues in the “wild-card” positions, even though these residues point away from the pocket domain (Jones et al., 1990; Palopoli et al., 2018; Singh et al., 2005). Analysis of our binding data and the sequence of the cellular proteins supports the hypothesis that bulky aromatic and perhaps to a lesser extent aliphatic sidechains in the wild-card spots confers higher affinity. For example, EID1, which has a glycine and aspartate in χ1 and χ2 (G,D), HDAC1 (A,E), and SMCA2 (T, E) all have affinities ~10 μM or greater. In contrast, ARID4A (V, H) and KMD5A (F, D) have affinities ~1 μM, and the viral E7 peptide (Y, Y) has submicromolar affinity.

The structures suggest two potential explanations for the increase in affinity conferred by bulky sidechains in the wild-card positions. In the Rb-bound E7 structure, the tyrosines at χ1 and χ2 are positioned to make stacking interactions (Figure 4A), demonstrating that large sidechains are able to contact each other for van der Waals interactions. In the case of p107-bound E7, however, the Y23 (χ1 position) is instead pointing toward and contacts V939 in p107, which is not conserved in Rb (M761) (Figure 4B). The Y25 (χ2 position) is still aligned in the p107 structure with its position in the Rb structure, sitting just above the backbone amide nitrogen and carbonyl of C24.

Notably, in the new structure of ARID4A here, H960 in the χ2 position also adopts the same x2 conformation, bringing it close to V958 (χ1 position in ARID4A) for van der Waals contacts and covering the C959 backbone nitrogen and carbonyl (Figure 4F). In all the solved LxCxE peptide structure, these cysteine backbone atoms form a bidentate hydrogen bond with the conserved N935 asparagine sidechain in the p107 pocket domain (N757 in Rb). The invariance of the χ2 sidechain orientation, also observed for D181 in EID1 (Figure 4D), suggests the additional role of shielding this critical hydrogen bond from water. We propose that the peptides with bulkier χ2 amino acids are more effective at shielding and bind with higher affinity. We found that mutation of N935A in p107 and N757A in Rb results in complete loss of detectable association by ITC for the high affinity E7 peptide, yet in control experiments to ensure the pocket structure is intact, the asparagine mutants still bind the E2F4 transactivation domain (Figures S3E-S3J). These data support the idea that the hydrogen bond interactions made by the asparagine are critical determinants required for LxCxE motif binding to the pocket domain.

#### Interactions C-terminal to the core LxCxE motif

It has been known from viral peptide structures and the LIN52 structure that a sidechain C-terminal to the LxCxE core motif fills a shallow hydrophobic pocket formed by the fourth and fifth cyclin fold helices and the intervening linker in the pocket domain (Guiley et al., 2015; Kim et al., 2001; Lee et al., 1998). In these previous structures, the residue filling that pocket is a leucine in the +2 position (or +2 methionine in the case of the SV40 T antigen). All the peptides in our binding studies have either a +2 or +3 hydrophobic residue, except notably SMCA2, mmBRG1, CycD1 and CycD2, which all have relatively weak affinity and harbor hydrophobic residues at +4 or +5 (Figure 1B). This comparison suggests that the +2/+3 hydrophobic interactions are important for high affinity, a conclusion also supported by mutagenesis of the HDAC1 sequence (Singh et al., 2005).

Comparison of previous structures and the new structures here with cellular LxCxE motifs reveals how the hydrophobic pocket can accommodate bulkier hydrophobics and a hydrophobic at the +3 position. This hydrophobic pocket is rather wide and shallow, and even the E7 (+2 Leu), LIN52 (+2 Leu) and EID1 (+2 Ile) peptides with similar amino acids in the +2 position fill the pocket with slightly different positions of the sidechain (Figure 5A). Another feature of the hydrophobic pocket is the pliability of M865 in p107, which forms the base of the pocket on the side near the fourth helix of the cyclin fold. In the HDAC1 structure, a bulkier phenylalanine in the +2 position (F420) is inserted deeper into the pocket, and the M865 sidechain accommodates by changing its (Figure 5B). An alternative M865 conformation is also observed in the ARID4A structure, which has a +3 Leu (L964). To accommodate the extra residue, the ARID4A main chain bulges out following the canonical E position (E961) and then approaches the hydrophobic pocket in p107 at a steeper angle. This results in the deeper insertion of L964 compared to the same side chain at the +2 position in other peptides, and L964 can be accommodated because M865 is again repositioned (Figure 5C).

**Figure 5.**
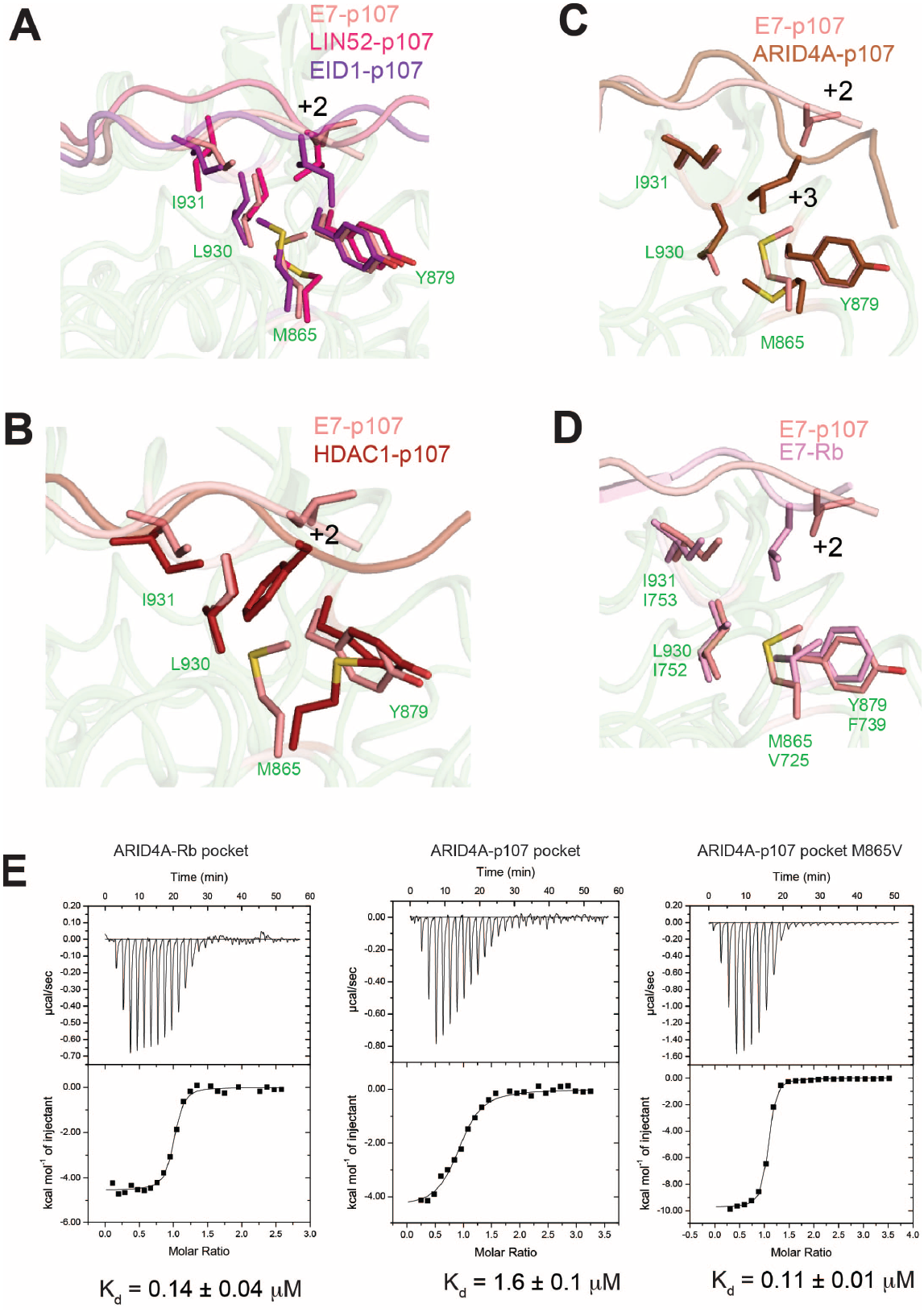
Flexibility in accommodating a +2/+3 hydrophobic sidechain. **(A)** Similar sidechains in the +2 position of E7, LIN52, and EID1 are oriented differently within the broad hydrophobic pocket of p107. **(B)** HDAC1 inserts its +2 phenylalanine deeper into the pocket, and M865 in p107 changes its position to avoid steric clash. **(C)** The +3 leucine of ARID4A is inserted deeper into the core relative to the E7 +2 leucine because of the angle of the main chain. M865 changes its position to accommodate the deeper leucine sidechain. **(D)** The V725 present in Rb at the same position as p107 M865 allows deeper insertion of the +2/+3 sidechain into the pocket without potential steric clash. (**E**) ITC measurements of ARID4A peptide binding to the indicated pocket domain construct. The reported K_d_ is the average of two replicates with the standard deviation reported as error.

The sequences C-terminal to the LxCxE motif in both the cellular and viral proteins have a number of acidic residues (Figure 1B), and truncation of these residues from peptides has been shown to weaken affinity (Chemes et al., 2011; Chemes et al., 2010; Palopoli et al., 2018; Singh et al., 2005). Addition of negative charge through phosphorylation has also been shown to increase affinity (Chemes et al., 2011; Chemes et al., 2010; Guiley et al., 2015; Palopoli et al., 2018; Pflum et al., 2001). In the case of LIN52, the structure shows the phosphoserine coordinated by a set of p107 pocket domain residues including two arginines (R880 and R869) that are conserved in Rb (Guiley et al., 2015). In contrast, we have found in these crystal structures that the C-terminal acidic residues either do not show ordered electron density (HDAC1), or they do not form any specific interactions with basic sidechains (ARID4A and EID1). We propose that this is due to the known diffuse long-range electrostatic interactions that drive the affinity (Chemes et al., 2011), which may be favored by a high degree of disorder in this region.

### Origin of higher affinity of Rb for several cellular LxCxE peptides

It is notable that the M865 position is not conserved in Rb, which has a valine at this position (V725) (Figure 5D). We hypothesize that the deeper pocket created by the shorter valine sidechain is a source of the higher affinity of several peptides for Rb (Figure 1B). We observed that the M865 orientation changes relative to the E7-p107 structure when p107 is bound with a peptide containing a bulky +2 phenylalanine (HDAC1, Figure 5B) or +3 hydrophobic (ARID4A, Figure 5C). This observation suggests that some peptides bind p107 weaker because they do not deeply bury the hydrophobic at the +2/+3 position or there is an energetic cost in repositioning M865. To test whether the presence of an Rb-like valine at the M865 in p107 confers higher affinity, we generated a p107 pocket domain construct with an M865V mutation and tested binding to the ARID4A peptide by ITC (Figure 5E). The affinities of ARID4A for wild-type Rb and p107 pocket domains were similar albeit slightly tighter than the affinities we measured by Alpha screen. Notably, data from both techniques demonstrate that ARID4A has a 10- to 30-fold greater affinity for Rb over p107 that we hypothesize arises from the requisite deep insertion of the +3 Leu into the hydrophobic pocket with the V725/M865 floor. We found in the ITC assay that the M865V p107 mutant binds the ARID4A peptide with ~15-fold greater affinity compared to wildtype p107 and with an affinity that is similar to its affinity for Rb.

The second sequence difference that may account for why some LxCxE motif peptides bind stronger to Rb is the presence of M761 in Rb compared to V939 in p107 at the same site. Here, the longer M761 sidechain in can make closer van der Waals contacts with the canonical L and χ1 sidechains in the motif as well as the peptide backbone (Figure 4). We found that a V939M p107 pocket domain mutant binds the ARID4A peptide with ~5-fold tighter affinity than wild-type p107, supporting the importance of the sidechain at this position (Figure S3K). However, it is not clear in this case why binding of some peptides, like ARID4A, are more affected by this substitution compared to other peptides (e.g. E7), which have similar affinity for Rb and p107. Another contributing factor may be that the longer M761 sidechain in Rb pushes the χ1 sidechain position toward the χ2 sidechain, which better covers the N757 hydrogen bonds (compare Figures 4A and 4B). This distinction may explain why the same Y23 at the χ1 position in E7 adopts a different orientation when bound to Rb or p107. The orientation of the χ1 phenylalanine in SV40 when bound to Rb similarly points toward the χ2 sidechain (Figure 4C). Together these observations suggest that differences in χ1 and χ2 sidechain rotamer conformations may be imparted by the M761/V939 substitution in Rb/p107.

### An engineered high affinity LxCxE-motif promotes association of HDAC1 with pocket proteins

We next introduced mutations into the HDAC1 sequence in order to engineer a higher affinity sequence. Based on our structural analysis and binding data (Singh et al., 2005), we chose mutations at the −1 position (R413D) and the wild card positions (A415Y, E417Y) that mimic the viral E7 LXCXE motif. We found using ITC that whereas the titration of p107 with the wild-type HDAC1 peptide yields poor signals and barely saturates using our conditions (we estimate K_d_ > 10 μM), the HDAC1 triple (3X) mutant peptide has affinity K_d_ = 105 ± 5 nM (Figure 6A). The HDAC1 mutant peptide also binds Rb tighter (Figure S3L), and we similarly observed that mutation of the wild-card residues in EID1 results in higher affinity for both p107 and Rb (Figures S3M-S3N). We determined the crystal structure of the HDAC1 3X mutant peptide bound to p107 and found that the tyrosines added to the wild-card positions are oriented similar to their position in the E7 peptide structure (Figure 6B). Y417 covers the backbone of C416 and Y415 is rotated toward V939. The D413 added to the −1 position makes the hydrogen bond interaction with the Y415 in the χ2 position observed in the Rb-bound E7 structure (Figure 6B). Overall, the conformation of the HDAC1 mutant peptide resembles more the structure of E7, which is consistent with the higher observed affinity and supports the conclusion that the orientation of bulky wild card residues is an important source of high affinity. To test the effects of a high affinity HDAC1 interaction in cells, we expressed Flag-tagged wild-type and mutant HDAC1 in HCT116 cells and assayed co-immunoprecipitation of endogenous Rb and p107. We tested the R413D, A415Y, and E417Y mutations individually and in combination. We performed the experiment in cells with either palbociclib or nocodazole treatment, which arrests cells in G1 with pocket proteins hypophosphorylated or in mitosis with pocket proteins hyperphosphorylated, respectively. We found that the triple mutant in particular more robustly associates with HDAC1 with Rb and p107 when cells are arrested in G1 (Figure 6C). In the nocodazole treated cells, we do not see a comparable increase in phosphorylated Rb or total p107 binding to the high-affinity HDAC1 mutant, suggesting that the effects of phosphorylation are still sufficient to dissociate pocket protein from the LxCxE motif. We note that we still observe an increase in band intensity when probing for total Rb in the immunoprecipitation from nocodazole cells, which, considering the result with the phosphospecific antibody, is most likely due to the presence of residual hypophosphorylated Rb.

**Figure 6.**
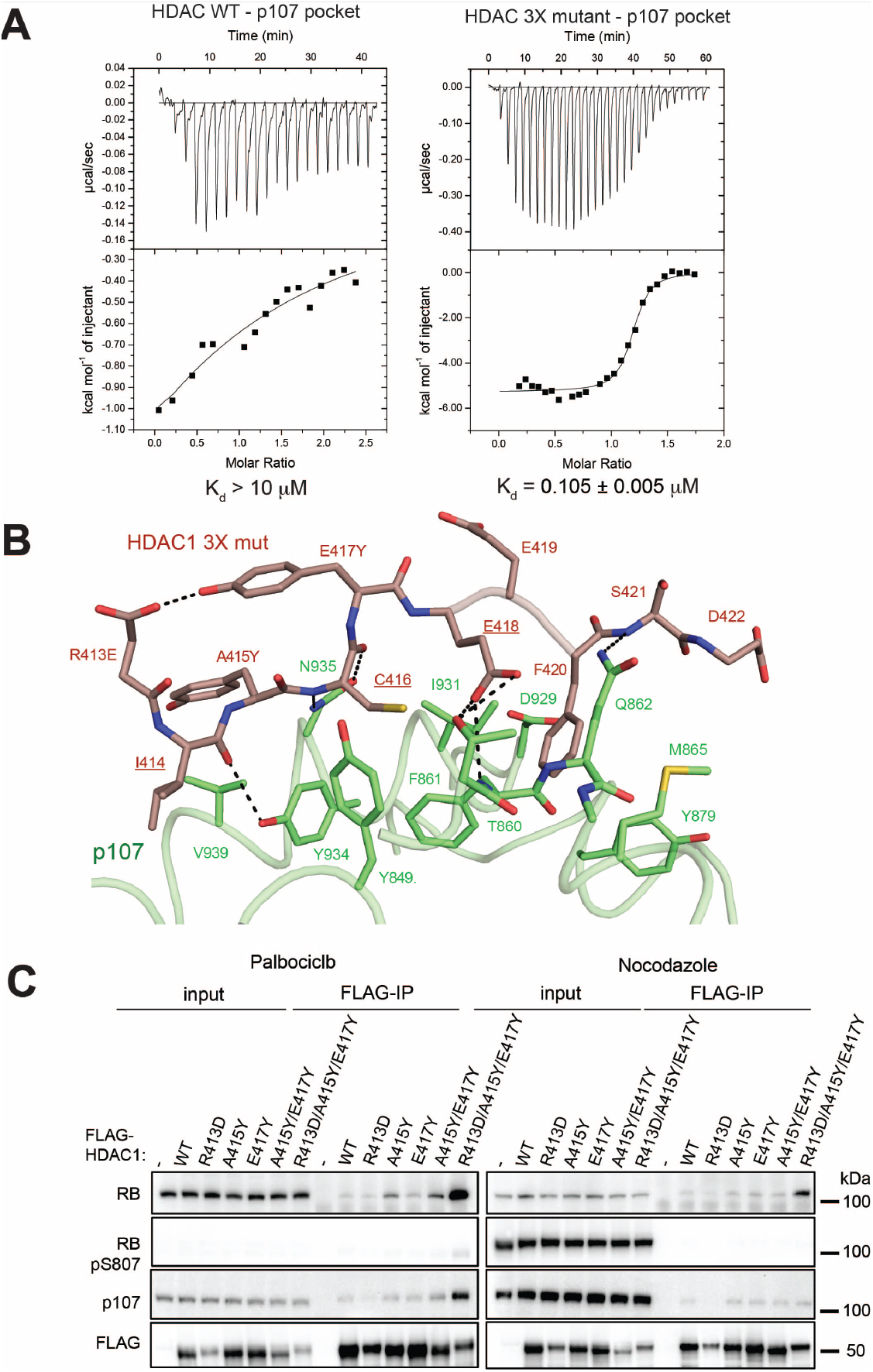
Mutation of the wild card residues and −1 acidic residue result in a high affinity HDAC1 interaction. (A) ITC data of HDAC1 3X mutant binding to the p107 pocket domain. (B) Crystal structure of HDAC3X bound to the pocket domain of p107. Mutations mimicking (D413, Y415 and Y417) the HPV16 E7 peptide were introduced into the HDAC1 LXCXE motif and show similar orientation as in the E7 peptide structures. (C) HCT116 cells transfected with the indicated HDAC1 construct were treated with drug for 24hrs prior to lysis, and immunoprecipitation was performed with an anti-Flag antibody.

## Discussion

While it has been shown that many cellular and viral proteins bind to Rb pocket proteins utilizing an LxCxE motif, there has been little systematic analysis of what sequence and structural features are critical for determining affinity. Here we present biochemical and structural data that further enhance our understanding of these determinants, particularly in the context of transcriptional corepressor protein interactions. Our data supports the hypothesis that a +2/+3 hydrophobic and bulky wild-card residues are important determinants of affinity and specificity and are often poorly optimized in the cellular proteins. Our results also provide structural explanations for several of these effects, for example, the protection of the N935 hydrogen bonds by bulky sidechains in the wild-card positions and the capacity of the +2/+3 sidechain to be deeply buried in a hydrophobic pocket. These design principles could be used in future efforts to synthesize viral-pocket protein inhibitors (Fera et al., 2012).

It remains an interesting question why viral proteins such as E7 have evolved to bind pocket proteins with high affinity, whereas cellular protein LxCxE motifs are often less optimized. In the case of viral proteins, very tight affinity may be required to target the pocket proteins early following infection or in differentiated epithelia, when viral effector protein concentrations are low (Banerjee et al., 2006; Crisostomo et al., 2019; Dowhanick et al., 1995). For cellular proteins, one possibility is that a weaker affinity may be needed for proper regulation of pocket protein interactions with functional partners. For example, the interactions of co-repressor proteins with Rb occur when Rb is hypohosphorylated, which is during G0/G1, and evidence suggests that specific phosphorylation events induce conformational changes that can disrupt binding to the LxCxE cleft (Burke et al., 2012; Ferreira et al., 1998; Harbour et al., 1999; Lai et al., 1999; Rubin, 2013). The idea that weak binding of an LxCxE motif better facilitates regulation has also been observed in the case of LIN52 binding to p107. The substitution of serine at the canonical cysteine position reduces binding, but phosphorylation of a C-terminal serine makes the association tighter (Guiley et al., 2015). We considered that cellular protein affinities are tuned specifically to bind pocket proteins only when they are in a hypophosphorylated state in the early cell cycle. However, we found that Rb and p107 phosphorylation is sufficient to dissociate the LxCxE-containing HDAC1 protein, even when the sequence is optimized to bind with high affinity similar to viral proteins. An alternative requirement for weak binding may be that it is needed for pocket proteins to form diverse complexes with a large pool of potential interacting partners, which can be switched through phosphoregulation or other molecular signaling events. It should also be noted that short linear interacting motifs are commonly found to have micromolar affinity, as they often make multivalent interactions and utilize cooperativity for signaling (Gibson, 2009; Kumar et al., 2021). In fact, CycD, HDAC1, and EID1 are all known to make multivalent interactions with Rb proteins (Hassler et al., 2007; MacLellan et al., 2000; Magnaghi-Jaulin et al., 1998; Miyake et al., 2000; Topacio et al., 2019).

A longstanding question about the pocket protein family is why Rb specifically has more potent tumor suppressor activity and is more often found mutated in cancer. One hypothesis is that subtle sequence differences between Rb, p107, and p130 account for different protein interactions that may mediate Rb-specific functions in suppressing growth or maintaining genomic integrity. Indeed, Rb has shown to specifically bind the activator E2Fs with high affinity, while only p107 and p130 are capable of binding LIN52 to assemble the DREAM complex (Guiley et al., 2015; Liban et al., 2017; Liban et al., 2016; Litovchick et al., 2007). Given the overall similarity of residues that form the LxCxE-binding cleft and observations of pocket protein associations with similar viral and cellular proteins containing the motif, it has been assumed that LxCxE-binding is a conserved function. In contrast, our results here suggest that Rb makes tighter interactions with several LxCxE peptides and that this higher affinity results from the greater ability of Rb to accommodate deep insertion of the +2/+3 hydrophobic into the pocket core. It is possible that these LxCxE-binding differences also contribute to differences in tumor suppressor potency. Notably, mutation of the LxCxE-cleft promotes tumorigenesis *in vivo* and has been linked to defects in genome stability and chromatin condensation (Coschi et al., 2010). Additional study of functional differences between pocket proteins that arise from different LxCxE-interactions, for example with transcriptional co-repressors, may reveal other key mechanisms of Rb tumor suppression.

## Materials and Methods

### Overexpression and Purification of p107 and Rb pocket domains

A GST fusion protein of the p107ΔL pocket domain (E389–Q972, ΔT600-779, Δ888-923) was expressed and purified from BL21(DE3) *E. coli* cells as previously described (Guiley et al., 2015). The culture, induced with 1 mM IPTG, was further grown for 15 hours at 18°C. Cells were harvested by centrifugation at 4000 rpm for 20 mins at 4°C. The cell pellets were lysed, and GS4B affinity chromatography was performed in 20 mM Tris (pH 8.0), 250 mM NaCl, 5 % Glycerol, 5 mM DTT and 1 mM PMSF. The GS4B purified protein was then subjected to GST-TEV protease cleavage and dialyzed overnight in 25 mM Tris, 250 mM NaCl, 5 mM DTT. The protein was then passed over GS4B affinity resin again to remove free GST and GST-TEV. Finally, the protein samples were concentrated and injected into a Superdex 75 Prep Grade column (GE Healthcare) in a buffer containing 20 mM Tris (pH 8.0), 500 mM NaCl, 5 mM DTT. For ITC, a similar construct of the human Rb pocket domain lacking the single internal loop (residues 380-787; Δ582-642) was expressed and purified using the same protocol. For the Alpha Screen measurements, His-tagged Rb pocket domain was produced as previously described (Chemes et al., 2010).

### Crystallization, data collection, structure determination, and model refinement

p107ΔL protein was crystallized following size-exclusion chromatography in complex with HDAC1, HDAC3X, ARI4DA and EID1 peptides by the hanging drop method at 4 °C. Peptides were added in twofold molar excess to 20 mg/mL p107ΔL and incubated for 30 min on ice before the crystallization. The reservoir crystallization solution consisted of 100 mM MES (pH 6.5), 1.6 M (NH4)2SO4, and 4% PEG 400. Crystals were flash frozen in the reservoir solution with 10 % glycerol. Data were collected at the Advanced Photon Source Beamline 23-D at 100K. Diffraction data from the ARI4DA-p107ΔL crystals were indexed, integrated, and scaled using XDS (Kabsch, 2010). The data from the other peptide-p107ΔL complex crystals were processed by using XIA2 (Winter, 2010) and AIMLESS (Evans, 2006) in the CCP4 program suite (Winn et al., 2011). ARI4DA and HDAC1-3X crystallized with p107ΔL in P1, and EID1 and HDAC1 crystallized with p107ΔL in C2_1_ space group. Structure determination of these complexes was achieved by using molecular replacement with PDB ID: 4YOS (LIN52-p107ΔL) as a search model (Vagin and Teplyakov, 2010). Alternative cycles of model building by COOT (Emsley et al., 2010) and refinement with Phenix (Afonine et al., 2010) were applied to complete the structures (Table S2).

### Peptide synthesis

Peptides for the AlphaScreen assay were provided by Belyntic GmbH. The peptides were purified by Belyntic’s Peptide Easy Clean (PEC) technology using a reductively cleavable linker system (PEC linker RC+) following the solid phase peptide synthesis (SPPS) adapted to plate format using an automated parallel peptide synthesizer (Intavis MultiPep RSi) (Zitterbart et al., 2021). The purified peptides were obtained as TFA salts in solid form in a 96 well collection plate. Identity and purity of the peptides was confirmed by UPLC-ESI-MS. For all the peptides except atCCD1, atCCD2, and the HPV16E7, a tyrosine was added to the N-terminus of the sequence shown in Figure 1B in order to quantify peptide concentration. An N-terminal alanine was added to atCCD21 to maintain five residues before the LxCxE motif and to HPV16E7 to replace the natural glutamine, and the C-terminal leucine in SMCA2 was replaced with a phenylalanine.

#### Alpha Screen assay for determination of binding affinities

An AlphaScreen™ assay was used to measure the inhibition of the interaction between a biotinylated E7-peptide (Biotin-QPETTDLYSYEQLNDS) and the human Rb or p107 pocket domains produced with an N-terminal 6xHis-tag (Rb) or an N-terminal GST-tag (p107). The E7 sequence was synthesized as a 16-mer peptide with C→S replacement and functionalized on the N-terminus with biotin. Binding of the biotin-E7 peptide to His-tagged Rb or GST-tagged p107 was detected with Streptavidin-coated donor beads and nickel chelate- or glutathione-coated acceptor beads. Concentration-response curves of inhibiting peptides were generated from 11-fold 1:3 serial dilutions starting at 100 μM with 6 nM His-Rb or GST-107, 6 nM biotin-E7 peptide and equal concentrations of 5 μg/mL acceptor and donor beads in 15 μl final assay volume. The assay was performed in white 384-well ProxiPlates and PBS supplemented with 2 mM DTT, 0.05% Tween 20 and 0.1% BSA as buffer. The AlphaScreen signal was measured in an Envision plate reader (Perkin Elmer, United States of America). Equilibrium dissociation constants from the AlphaScreen assay were obtained by fitting the normalized data and considering the binding equilibrium of both the biotin-E7 peptide and the competitor peptides to Rb or p107 according to a previously described model (Kuzmic et al., 1992). Data fitting was performed using pro Fit 7 (Quantumsoft, Zurich, Switzerland).

### Isothermal titration calorimetry (ITC)

Prior to ITC measurements, peptides and proteins were dialyzed overnight against a buffer containing 20 mM Tris (pH 8.0), 150 mM NaCl, and 2 mM BME at 4°C. Equilibrium dissociation constants were obtained at 4°C using a MicroCal VP-ITC system (GE Healthcare). Peptide concentrations between 300-750 μM in the syringe were injected into 25–35 μM pocket domain in the cell. A single binding site model was employed to determine the equilibrium dissociation constants. In each titration, the stoichiometry of peptide binding to pocket proteins was measured to be close to 1:1 (n ~ 0.9-1.1).

### Co-immunoprecipitation

Plasmids for the expression of HDAC1 mutants in mammalian cells were generated by site-directed mutagenesis following the Stratagene QuikChange mutagenesis protocol. The plasmid expressing FLAG-tagged wild-type human HDAC1 was a gift from Eric Verdin (Addgene Plasmid # 13820). HCT116 cells were transfected in 10 cm plates with 8 μg plasmid and 40 μl PEI. 24 h after transfection, cells were treated with either 1 μM Palbociclib or 1 μl Nocodazole for additional 24 h. Cells were lysed with IP buffer containing (Tris pH 8.0, 0.2% Triton X, protease inhibitors (EDTA-free protease inhibitor tables, Pierce)) supplemented with 300 mM NaCl by incubation for 10 min on ice followed by direct sonication (5 times for 1 s). Lysates were cleared by centrifugation (4 °C, 10 min, 20,000 xg) and diluted with IP buffer to an NaCl concentration of 150mM. 4mg protein in 2ml IP buffer were incubated with 40μl Anti-DYKDDDDK Magnetic Agarose (Pierce) on a rotator for 30min at 4°C. Beads were washed 5x with IP buffer/150 mM NaCl and bound proteins were eluted with 50 μl 1x Laemmli buffer at 95°C. SDS-PAGE. Western blots were performed following standard protocols. The following antibodies were applied to detect RB, p107, and HDAC-FLAG: RB-C2 (Santa Cruz Biotech, sc-74562, RRID:AB_2177334), RB phospho S807 (Abcam, ab47762, RRID:AB_882296), p107-D3P3C (Cell Signaling Technology, #89798, RRID:AB_2800144), and Anti-OctA-Probe H-5 (Santa Cruz Biotech, sc-166355, RRID:AB_2017593).

## Acknowledgements

This work was supported by grants from the National Institutes of Health to S.M.R. (R01GM124148 and R01GM127707), Ministerio de Ciencia, Tecnología e Innovación Productiva and Deutscher Akademischer Austauschdienst (MinCyT-DAAD) to L.B.C. and T.J.G. (CyCMotif DA/16/05), and Agencia Nacional de Promocion Científica y Tecnológica (ANPCyT) Grant #PICT 2017/1924 to L.B.C. This research has received funding from the European Union’s Horizon 2020 research and innovation program to L.B.C. and T.J.G. under the Marie Skłodowska-Curie grant agreement no. 778247 (IDPfun). L.B.C. is a National Research Council Investigator (CONICET, Argentina), and L.A was supported by a CONICET postdoctoral fellowship.

## Author Contribution Statement

Conceptualization and experimental design: L.B.C. and S.M.R. Sequence analysis: L.B.C. and T.J.G. Performance and experimental design of crystallization and ITC assays: S.P., S.M.F., and S.T. Preparation and design of peptide sequences and affinity data analysis: L.A. Peptide Synthesis: S.L. Peptide measurement using AlphaScreen assay: J.L and P.S. Cellular assay experimental design and performance: G.A.M. Manuscript drafting: S.P., S.M.R and L.B.C. Manuscript editing: S.P., L.A, G.A.M., T.J.G, L.B.C. and S.M.R.

---

**Table S1.**
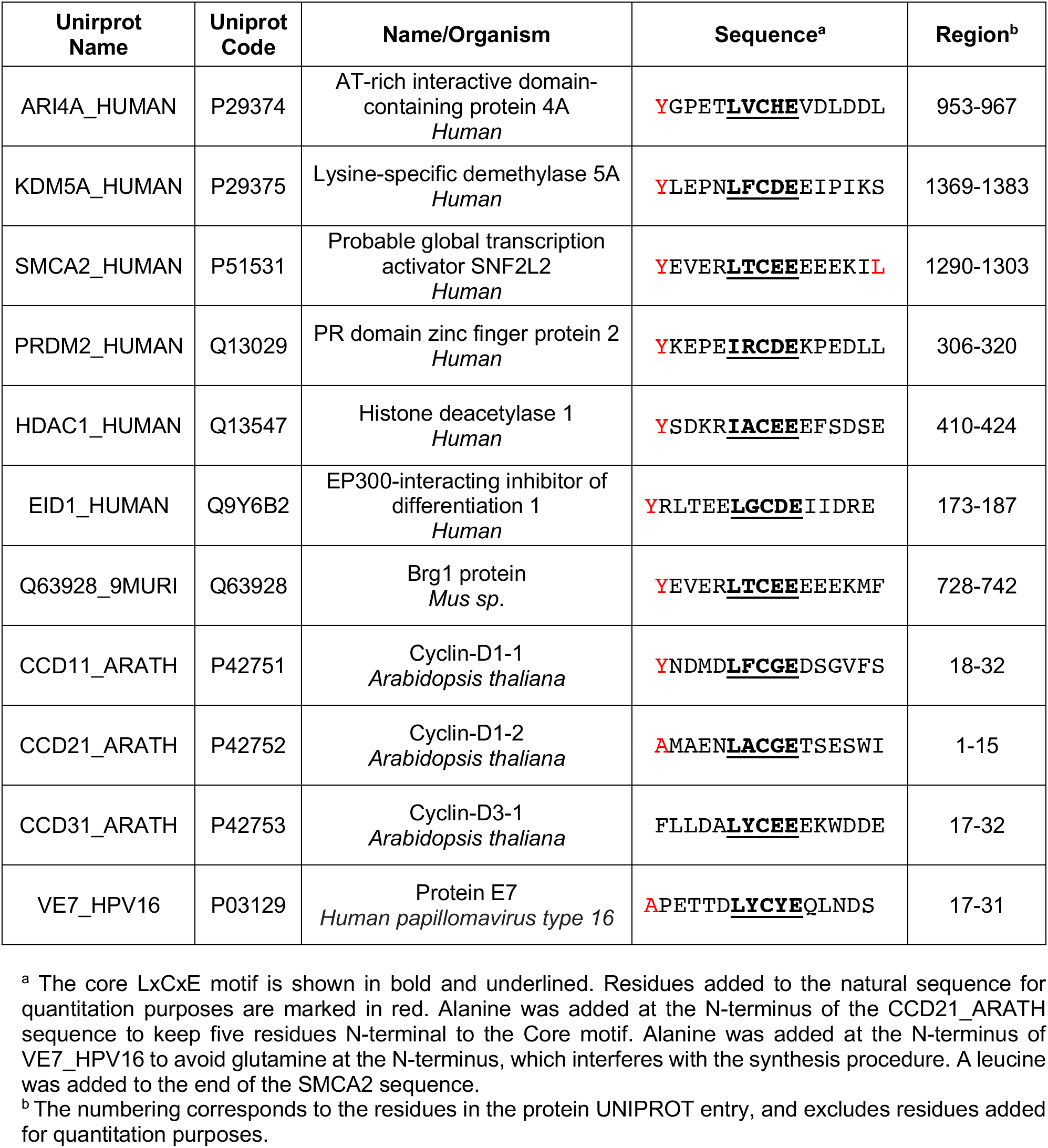
Uniprot sequence information for cellular and viral LxCxE peptides.

**Table S2.**
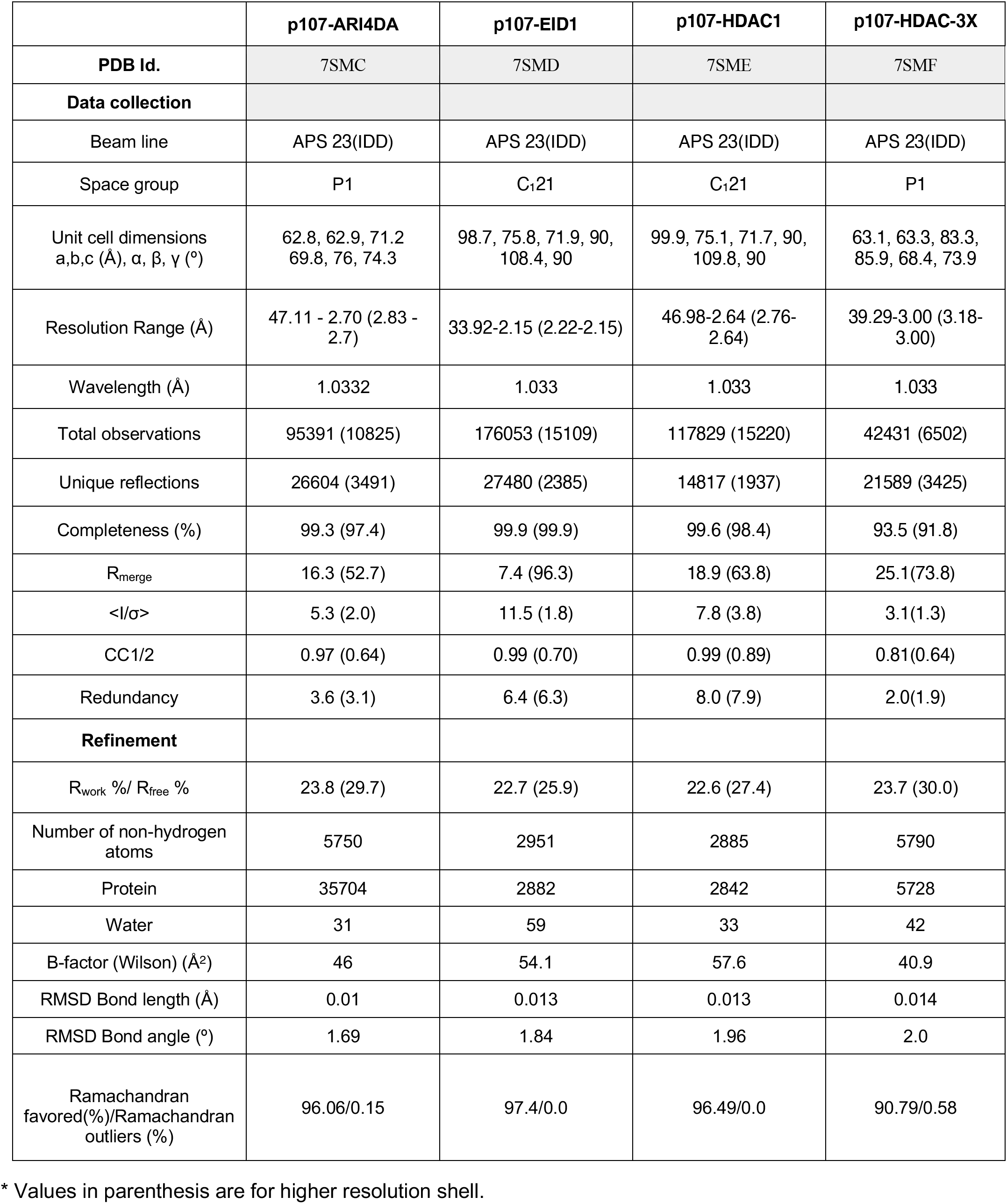

**Figure S1:**
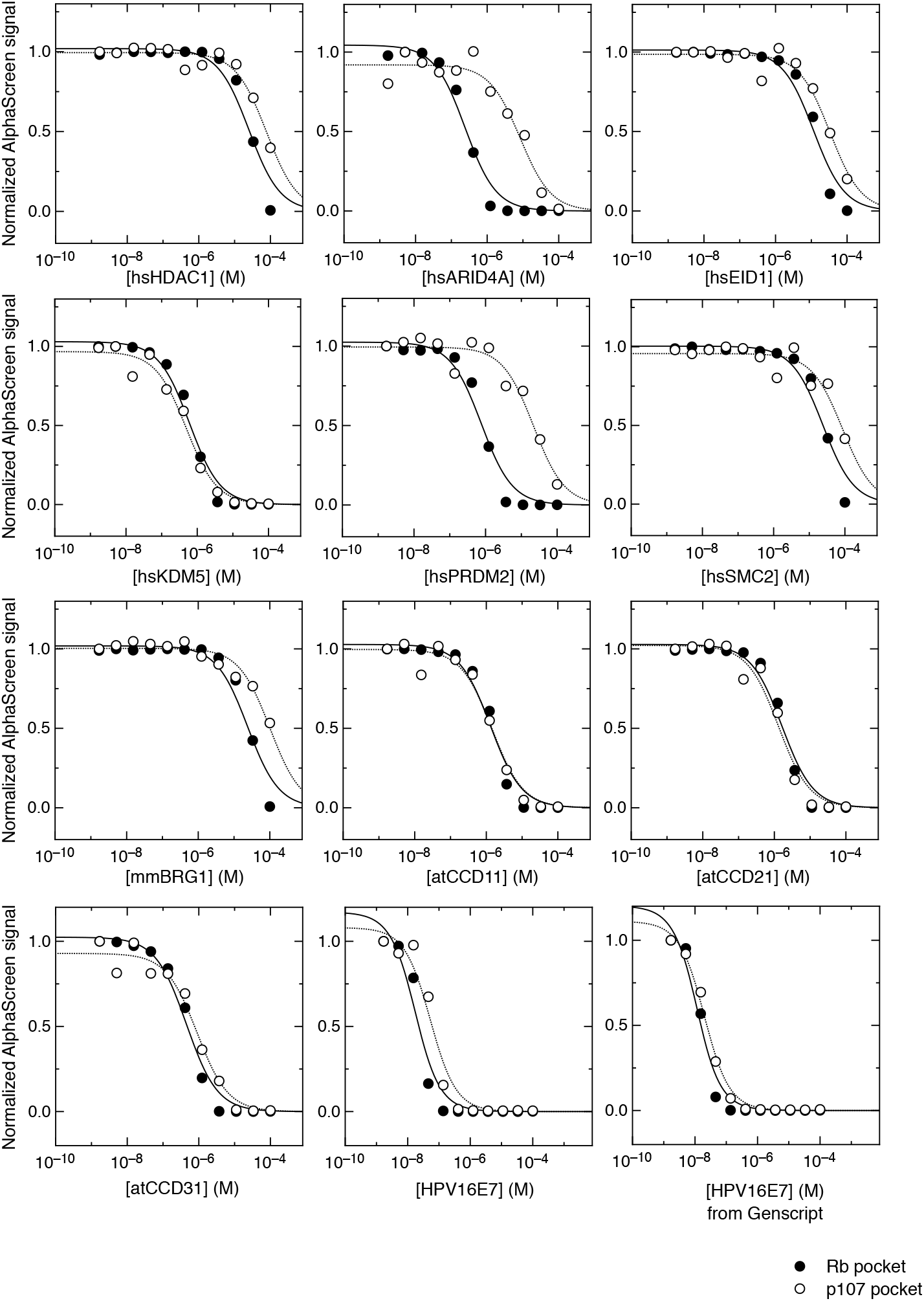
Alpha Screen affinity measurements of LxCxE peptides for the Rb and p107 pocket domains. Competition of a preformed complex of 6nM His-Rb or GST-107 and 6nM biotin-E7 peptide with different LxCxE peptides. Competitor peptides were: hsHDAC1, hsARID4A, hsEID1, hsKDM5A, hsPRMD2, hsSMC2, mmBRG1, atCCD11, atCCD21, atCCD31 and HPV16E7. Black circles: Competition of [biotinE7-Rb] complex. Open circles: Competition of [biotinE7-p107] complex. Affinity for the HPV16E7 peptide from Belyntic was (K_d_ = 0.014 ± 0.005 for Rb and K_d_ = 0.050 ± 0.010 for p107) and that for the HPV16 E7 peptide from Genscript was (K_d_ = 0.008 ± 0.002 for Rb and K_d_ = 0.015 ± 0.020 for p107).

**Figure S2:**
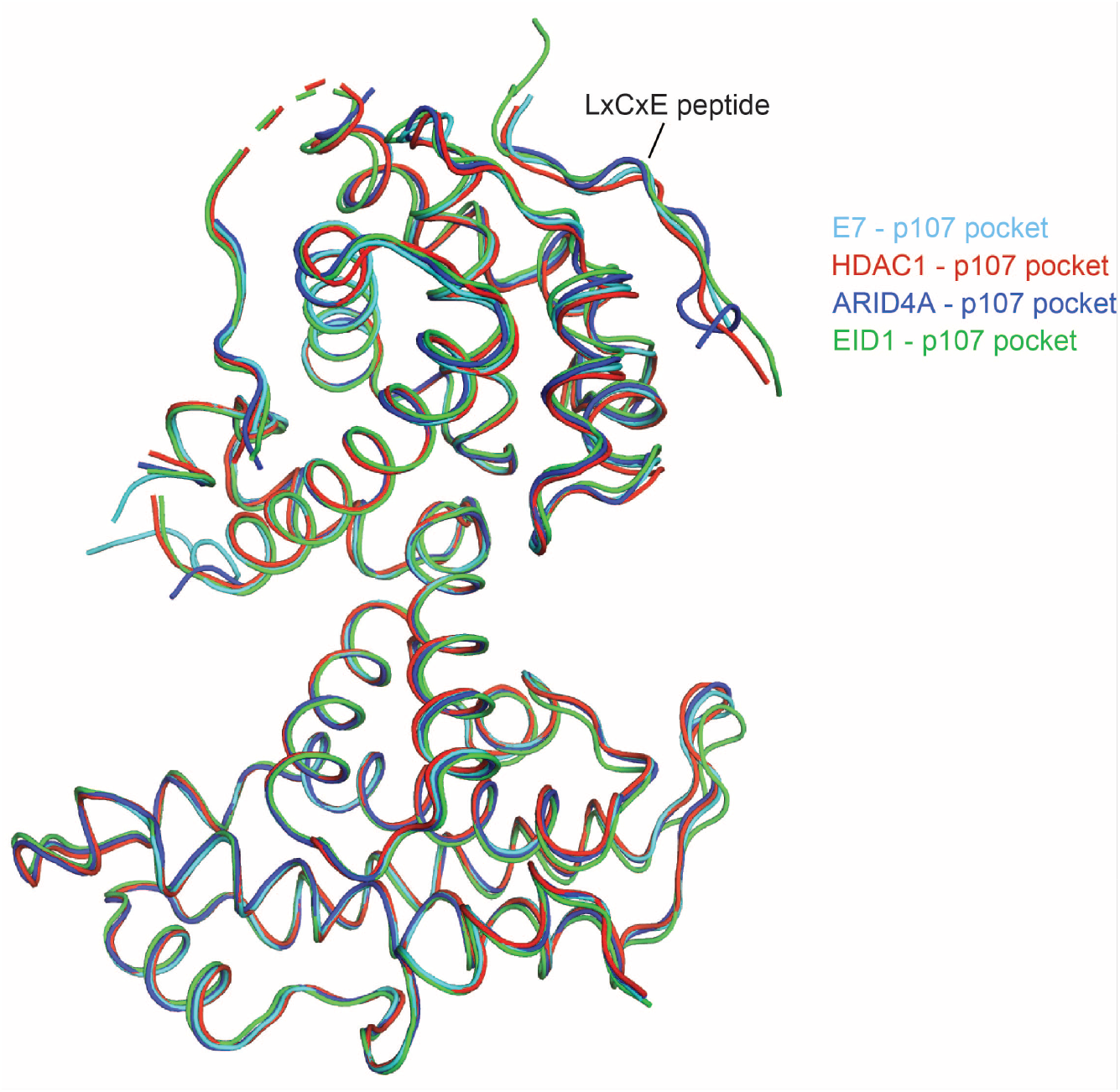
Superposition of the p107 pocket domain from structures with different bound LxCxE peptides. The structures with HDAC1 (red), ARI4DA (blue), and EID1 (green) bound have root mean square deviations (RMSD) of 0.32 Å, 0.26 Å, and 0.49 Å for Ca positions compared to p107 in the E7 complex structure (cyan).

**Figure S3:**
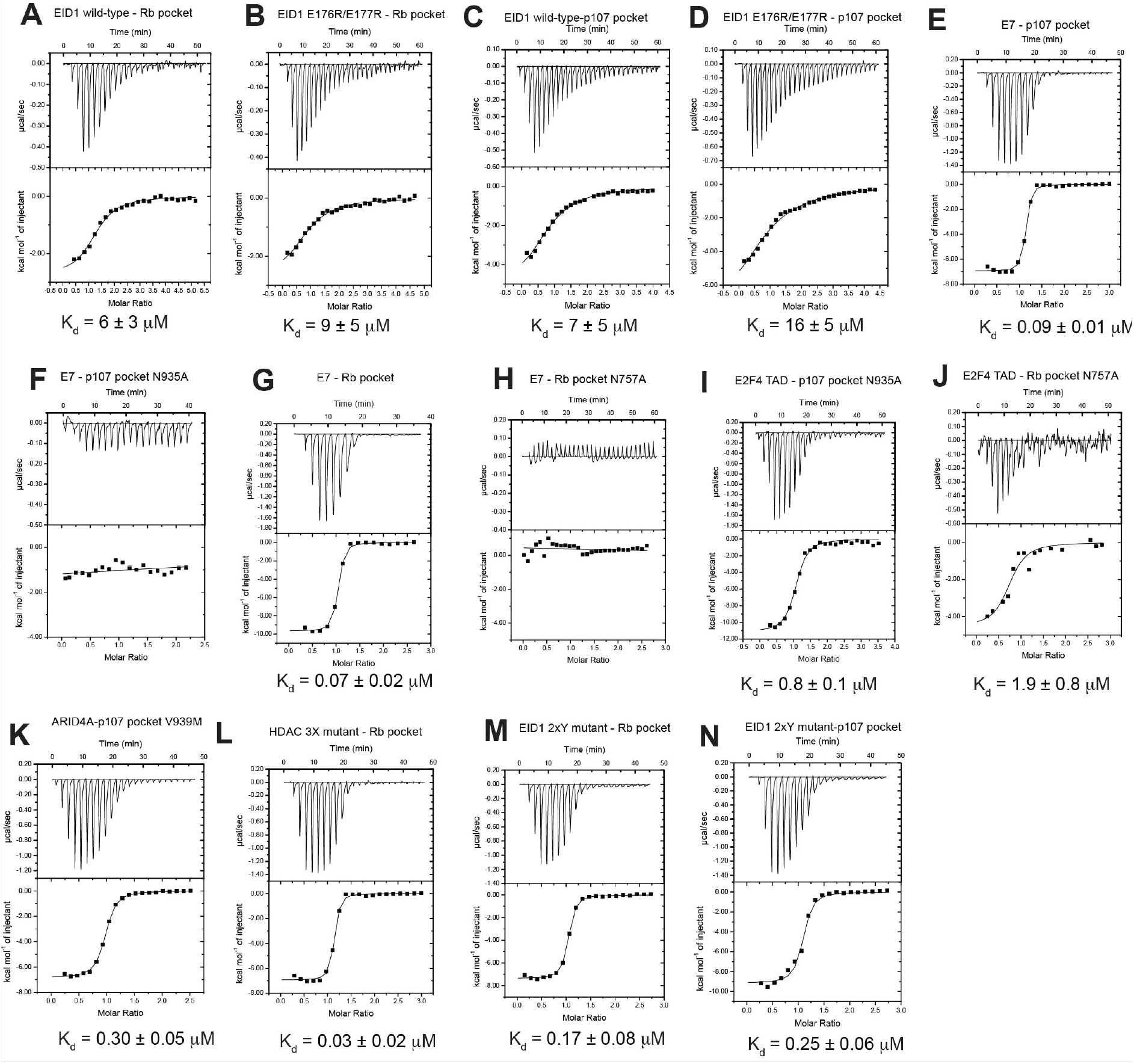
Supplemental ITC binding data. Representative ITC titration curves for the indicated peptide-protein interaction. The reported K_d_ values are averages from 2-3 replicates with standard deviations shown as the error. The E2F4 transactivation domain (TAD) peptide contains the sequence HDYIYNLDESEGVCDLFD (residues 390-407). The affinities measured for the E2F4 TAD association with the mutant pocket domains were similar but 4-5 fold lower than that previously reported (Liban et al., 2016).

## Notes

### Competing Interest Statement

The authors have declared no competing interest.

